# The SARS-CoV-2 spike protein alters barrier function in 2D static and 3D microfluidic in vitro models of the human blood–brain barrier

**DOI:** 10.1101/2020.06.15.150912

**Authors:** Tetyana P. Buzhdygan, Brandon J. DeOre, Abigail Baldwin-Leclair, Hannah McGary, Roshanak Razmpour, Peter A. Galie, Raghava Potula, Allison M. Andrews, Servio H. Ramirez

## Abstract

As researchers across the globe have focused their attention on understanding SARS-CoV-2, the picture that is emerging is that of a virus that has serious effects on the vasculature in multiple organ systems including the cerebral vasculature. Observed effects on the central nervous system includes neurological symptoms (headache, nausea, dizziness), fatal microclot formation and in rare cases encephalitis. However, our understanding of how the virus causes these mild to severe neurological symptoms and how the cerebral vasculature is impacted remains unclear. Thus, the results presented in this report explored whether deleterious outcomes from the SARS-COV-2 viral spike protein on primary human brain microvascular endothelial cells (hBMVECs) could be observed. First, using postmortem brain tissue, we show that the angiotensin converting enzyme 2 or ACE2 (a known binding target for the SARS-CoV-2 spike protein), is expressed throughout various caliber vessels in the frontal cortex. Additionally, ACE2 was also detectable in primary human brain microvascular endothelial (hBMVEC) maintained under cell culture conditions. Analysis for cell viability revealed that neither the S1, S2 or a truncated form of the S1 containing only the RBD had minimal effects on hBMVEC viability within a 48hr exposure window. However, when the viral spike proteins were introduced into model systems that recapitulate the essential features of the Blood-Brain Barrier (BBB), breach to the barrier was evident in various degrees depending on the spike protein subunit tested. Key to our findings is the demonstration that S1 promotes loss of barrier integrity in an advanced 3D microfluid model of the human BBB, a platform that most closely resembles the human physiological conditions at this CNS interface. Subsequent analysis also showed the ability for SARS-CoV-2 spike proteins to trigger a pro-inflammatory response on brain endothelial cells that may contribute to an altered state of BBB function. Together, these results are the first to show the direct impact that the SARS-CoV-2 spike protein could have on brain endothelial cells; thereby offering a plausible explanation for the neurological consequences seen in COVID-19 patients.

## INTRODUCTION

Coronavirus disease 2019 (COVID-19) is a rapidly emerging public health crises threatening human health. Initially reported as pneumonia of unknown origins from Wuhan, Hubei Province, China. COVID-19 infection is caused by severe acute respiratory syndrome corona virus 2 (SARS-CoV-2) a novel human β-coronavirus that shares highly homological sequence with SARS-CoV^1,2^. Infection is primarily transmitted by respiratory droplets and from human to human contact with a median incubation period of approximately 5 days^3^.

The clinical spectrum of COVID-19 varies from asymptomatic, mild to moderate self-limiting disease in the majority of cases^4-6^. However, severe and fatal consequence can occur in some patients with comorbidities such as cardiovascular/pulmonary disease and diabetes^7,8^. The most common symptomology in patients includes fever, dry cough, fatigue, diarrhea, alteration in taste/smell, conjunctivitis, and pneumonia^3,9^. Patients with severe respiratory infection can progress to acute respiratory distress syndrome (ARDS) and multiple organ failure^10^. Severity of disease in COVID-19 patients is associated with immune system dysregulation, such as lymphopenia and inflammatory cytokine storm. Among the other distinctive features of COVID-19 are the elevated D-dimer, C-reactive protein, procalcitonin, LDH, ferritin, LFTs, reduced CD4+/CD8+ T cells, predictive of mortality^11-13^.

The pulmonary system is principally involved in fatal COVID-19 cases. Histopathological observation from several autopsy studies have described diffuse alveolar damage with necrosis of alveolar lining cells, pronounced reactive type II pneumocytes, linear intraalveolar fibrin deposition and hyaline membrane formation consistent with diffuse alveolar damage^14,15^. Additionally, evidence suggest broad tropism for SARS-CoV-2 in the kidneys, heart, large intestines, spleen, and liver^16^. Nevertheless, despite widespread interest in the pathophysiology of the disease, much remains unknown about how SARS-CoV-2 affects the CNS. Neurological signs, such as headache, nausea, vomiting and impaired consciousness have been reported^17,18^ with COVID-19, intriguing the plausibility of SARS-CoV-2 to invade the central nervous system resulting in neurological diseases^19^. Other coronaviruses have been shown to be neurotropic, causing encephalitis in felines^20^, porcine^21^ and murine^22^. Additionally, SARS-CoV-1, the most closely related coronavirus, has been shown to infect the brain stem of both human and animals^23^ and the first case of meningitis/encephalitis has been reported in a patient associated with SARS-CoV-2^24^. Consequently, there is also growing evidence that SARS-CoV-2 has direct effects on the vasculature, specifically in the central nervous system. The virus binds to angiotensin converting enzyme 2 (ACE2)^25^, a cell surface carboxypeptidase part of the renin-angiotensin system (RAS) that is responsible for a host of functions in the cardiovascular system. Specifically, ACE2 catalyzes the degradation of angiotensin II to angiotensin fragment (1-7) (Ang-(1-7)), which is associated with vasodilation and a subsequent decrease in hypertension^26,27^. In fact, recent studies have found increased serum levels of angiotensin II in COVID-19 patients^28^. ACE2 is expressed on the vasculature^2^ and recent studies of COVID-19 patients have shown virus like particles in pulmonary^29^ and kidney^30^ endothelium. ACE2 is also expressed on the human cerebral vasculature^19,31^, which we also confirm herein. However, how the virus and engagement of ACE2 alters the blood-brain barrier and contributes to possible neuroinvasion remains an open question.

The present study aimed to identify whether SARS-CoV-2 spike proteins negatively affects brain endothelial cells. Here, it is shown that the primary cellular binding target for the S1 protein, ACE2, is present throughout the human cerebral vasculature. Functional outcomes were performed using primary human brain microvascular endothelial cells in both 2D (monolayer) and 3D (endothelialized cylindrical voids within hydrogels) for recapitulation of the BBB. With the above BBB models, the effect of SARS-COV-2 spike subunits on barrier integrity were determined. Evidence is provided that points to the negatively impact that the SARS-CoV-2 spike proteins have on barrier function that may be explained via pro-inflammatory responses of endothelial cells to these viral proteins. To the authors knowledge, this is the first evaluation for the effects of SARS-CoV-2 spike protein on the blood brain barrier.

## MATERIALS AND METHODS

### Reagents

Spike protein S1 subunit (RayBiotech, Cat No 230-01101), Spike protein S1 subunit Receptor Binding Domain sequence (RBD) (RayBiotech, Cat No 230-01102), and Spike protein S2 subunit (RayBiotech, Cat No 230-01103) were used for in vitro experiments. Other recombinant proteins (i.e TNFα) were purchased from R&D Systems (Minneapolis, MN, USA).

### Endothelial Cell Culture

All monolayer experiments used primary human brain endothelial cells derived from fetal brain tissue. Healthy tissue was provided (under informed consent) by the Laboratory of Developmental Biology (University of Washington, Seattle, WA) with approval granted by Temple University’s (Philadelphia, PA) Institutional Review Board and in full compliance by the National Institutes of Health’s (NIH) ethical guidelines. Fetal human brain microvascular endothelial cells (hBMVEC) were isolated as described ^32^. Cells were grown on rat-tail collagen I coated flasks (BD Biosciences) in EBM-2 medium supplemented with EGM-2MV SingleQuots (Lonza, Cat No CC-3156 and CC-4147) in an incubator set to 37 °C, 5 % CO_2_, and 100 % humidity.

### Immunohistochemistry and Imaging

Formalin-fixed paraffin-embedded human frontal cortex brain tissue blocks of both normal and diseased origin (mixed-type dementia, cardiovascular and pulmonary dysfunction) were procured from ProteoGenex, Inc. (Inglewood, CA), and serially sectioned at a thickness of 5 µm each. Glass slide-mounted sections were cleared, rehydrated and placed through heat-induced epitope retrieval (HIER) using Tris-EDTA buffer (pH 9.0) in preparation for immunohistochemical staining. HIER pre-treated sections were blocked for endogenous alkaline phosphatase activity and non-specific antibody binding using Bloxall (Vector Laboratories, SP-6000) and 2.5% Normal Horse Serum (Vector Laboratories, S-2012), respectively. The sections were subsequently incubated in rabbit anti-human ACE2 antibody (1:500, Abcam, ab15348) for 1 hr at RT. Positive antibody binding was detected using anti-rabbit IgG ImmPRESS-AP polymer reagent (Vector Laboratories, MP-5401) and visualized via a 10 min incubation in Vector Blue AP substrate (Vector Laboratories, SK-5300). Stained sections were dehydrated, cleared and permanently mounted with VectaMount (Vector Laboratories, H-5000) for subsequent bright field imaging.

For Imaging, all sections were scanned with the Aperio AT2 slide scanner (Leica Biosystems) and analyzed using Aperio ImageScope software (v12.3.2.8013). These cortical regions of brain tissue were examined in 4 mm by 4 mm regions of interest for expression of ACE2 on blood vessels (based on morphological appearance).

### 3D BBB model

Three-dimensional models of the blood-brain barrier (3D BBB) were fabricated by polymerizing hydrogels composed of 5 mg/mL type I collagen, 1 mg/mL hyaluronan, and 1 mg/mL Matrigel within microfabricated devices. The full method for this approach is described in a previous study^33^. Briefly, hydrogels were injected into the reservoir of the device and 180-µm needles coated in 0.1% BSA were inserted prior to polymerization to create two, parallel, and cylindrical voids within the gel. HCMEC/D3 were injected into one channel at a density of 10 million per mL (15 µL per channel). Channels were incubated for 10 minutes to ensure cell attachment then injected with cells again and inverted for 10 minutes to coat the opposing side to ensure full coverage. Following cell seeding, channels were exposed to 0.7 dyn/cm^2^ of steady shear stress for four days using a linear syringe pump (Kent Scientific) to establish barrier function. Following the four-day perfusion, vessels were perfused for two hours with 50 nM of SARS-CoV-2 S1 spike peptide (ProSci). Following exposure to the spike protein, vessels were either placed in fixative or prepared for permeability testing.

### Permeability Assays (Static)

To evaluate the paracellular permeability under static conditions, cells were seeded at the density of 10,000 cell per collagen I coated Transwel insert (pore size 0.4 μm, diameter 0.33 cm^2^, Corning) in the 200 μL of EGM2-MV medium. Basolateral chambers were filled with 500 μL of medium. Medium was changed every 3 days. After confluent monolayer was formed, hBMVEC monolayers were incubated with 10 ng/ml TNF-α, or 10nM SARS-CoV-2 spike protein subunit S1, spike protein subunit S2 or receptor binding domain (RBD) sequence of S1 (RayBiotech, Cat No 230-01101, 230-01103, 230-01102 correspondingly). 3kDa FITC-DEAE-conjugated dextran (Sigma) was added to the apical chamber to the final concentration of 1 mg/ml and 1 h later, fluorescence in the basolateral chambers was determined using a SpectraMax M5e (Molecular Devices). Percent permeability was calculated as the relative fluorescence of medium in the spike protein-treated vs untreated cells.

For permeability measurements in 3D, vessels were transferred to the stage of an inverted epifluorescent microscope enclosed by an environmental chamber set to 37C, 5% CO2, and 95% RH. The channels were perfused with 4-kDa dextran-FITC at a flow rate of 5 µL/min using a syringe pump for 10 minutes, while submerged within culture medium to ensure cell viability. This flow rate was selected to assure fully developed flow throughout the channel and to maintain consistency with previous work. Images were taken at 30s intervals for 10 minutes, and the diffusion coefficients were established using the following equation from previous work^34^.

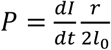

Where P is the permeability coefficient, dI/dt is the rate of change in fluorescence intensity outside the 3D vessel, r is the vessel radius, and I_0_ is the fluorescence intensity inside the 3D vessel.

### Cytotoxicity Assay

The LIVE/DEAD viability/cytotoxicity assay (Life Technologies, Cat No L3224) was used to evaluate SARS-CoV-2 spike proteins toxicity to hBMVEC cell. Briefly, hBMVEC cells were seeded on a sterile 96-well plate at 1 × 10^4^ cells per well and grown to confluency. Confluent cells were treated with 10nM SARS-CoV spike protein subunit S1, spike protein subunit S2 or receptor binding domain (RBD) sequence of S1 for 48 h. 200 μL of 1 μM calcein-AM and 5uM ethidium homodimer-1 were added to each well and incubated for 30 min at room temperature. Data was acquired at excitation and emission wavelengths of 495/515 nm for live cells and 528/617 nm for dead cells and normalized to control (untreated) cells.

### Flow Cytometry

Cells plated in 12-well dishes were grown to confluency and treated with 10nM of S1 full, RBD, S2 full or 100ng/mL of TNF-α, washed with calcium and magnesium free PBS and then detached with accutase for 1-2 min at 37°C. Cells were then pelleted by centrifugation at 1000 rpm for 5 min and re-suspended in fixation buffer (ebioscience/Thermo Fisher) for 30 min. Following fixation, cells were washed with flow cytometry buffer (5% FBS 0.1% sodium azide) and pelleted again. Cells were resuspended for 30 min in 100 μL of flow cytometry buffer and anti-ICAM-1 (Pe-Cy7, Biolegend) and anti-VCAM (APC, Biolegend) preconjugated primary antibodies. Cells were then washed, pelleted and resuspended in flow cytometry buffer for FACS analysis. 10,000 events per sample were acquired with a FACS BD Canto II flow cytometer (BD Biosciences) and data was then analyzed with FlowJo software (Tree Star, Ashland, OR, USA).

### Quantitative real time PCR

To examine the concentration of mRNA, total RNA was extracted using TRIzol and PureLink RNA extraction reagents (Invitrogen). cDNA was synthesized with 400 ng of RNA in 20 μL reaction mix using High Capacity cDNA Reverse Transcriptase kit (Applied Biosystems). qRT-PCR was perfomed using TaqMan Universal 2x Master Mix (Thermo Scientific) and human TIMP-1 (Hs01092512), MMP2 (Hs01548727), MMP3 (Hs00968305), MMP9 (Hs00957562), MMP12 (Hs00159178), IL1b (Hs01555410), IL6 (Hs00174131), CXCL10 (Hs00171042), CCL5 (Hs00982282) FAM-labeled probes. 18S rRNA (Cat No 4352930). was used as an internal control. Gene expression levels were analyzed using 2 ^-ΔΔCt^ algorithm.

### Protein electrophoresis and immunoblotting

Confluent cell monolayers were treated with SARS-CoV-2 spike proteins for 24 hrs and briefly rinsed with PBS. Whole cell lysate was prepared using RIPA buffer (EMD Milipore, Cat No 20-188) as per manufacturer’s protocol. Obtained fractions were subjected to 10% Bis-Tris polyacrylamide gel electrophoresis in MOPS buffer under denaturing conditions and transferred to a 0.45 μm PVDF membrane. Membrane was blocked with Odyssey blocking buffer in Phosphate-buffered saline (Li-Cor Biosciences, Cat No 927-40000) for 1 h at room temperature. Blocked protein blot was incubated with affinity-purified rabbit anti-ACE2 (1:1000, Abcam, Cat No 15348), and mouse anti-b-actin (1:5000, Sigma, Cat No A5441), in PBS with 0.05% Tween-20 and 10% Odyssey blocking buffer at 4°C overnight, followed by incubation with goat anti-rabbit IRDye 800CW IgG and goat anti-mouse IRDye 680RD secondary antibody in PBS (1:20,000) at room temperature for 1 h. Protein blot was visualized with Odyssey LiCor Imaging System. Band intensities were quantified using ImageJ software (NIH, Bethesda). Data is presented as relative intensity of ACE2 bands in SARS-CoV-2 spike proteins / untreated bands normalized to β-Actin.

### Statistical analysis

The experiments were independently performed multiple times (at least three times for all the data shown) to allow statistical analyses. Within each individual experimental set, primary cells from at least 3 donors were used, and every condition/per donor was evaluated in at least three replicates. The data collected was analyzed using Prism v6.0 (GraphPad Software, San Diego, CA). All the results are expressed as the mean ± SD with differences considered significant at *p* < 0.05.

### ACE2 expression in human cerebral vasculature and in primary human brain endothelial cells in culture

Previous reports on the structure and biochemical interactions of the SARS-CoV-2 with cellular protein targets have revealed that binding to the ACE2 membranous protein is a critical step for SARS-CoV-2 to enter target cells. To evaluate whether the Angiotensin-Converting Enzyme 2 (ACE-2), is expressed in the brain vasculature, immunostaining for the ACE2 protein was performed on postmortem brain tissues from normal and with neurodegenerative disease (mixed dementia). As shown in Figure 1, expression of ACE2 is clearly detected in the frontal cortical regions of the brain. ACE2 appears in small vessels such those that form capillary networks (Figure 1A, top left). Immunopositive staining for ACE2 is also observed is larger caliber vessels (Figure 1A, top left, bottom left and bottom right). Cross-sectioned vessels shows staining throughout the vessel wall and within the medial layers. In case of mix dementia, immunopositive ACE2 expression is found at higher levels in the brain parenchyma and in all types of vessels (Figure 1B). Thus, we corroborate previous findings of ACE2 expression in the brain and further demonstrate that all vessel calibers express the protein. Furthermore, ACE2 levels may be altered in the cerebral vasculature as a result of chronic neuroinflammation associated with neurodegeneration.

**Figure 1.**
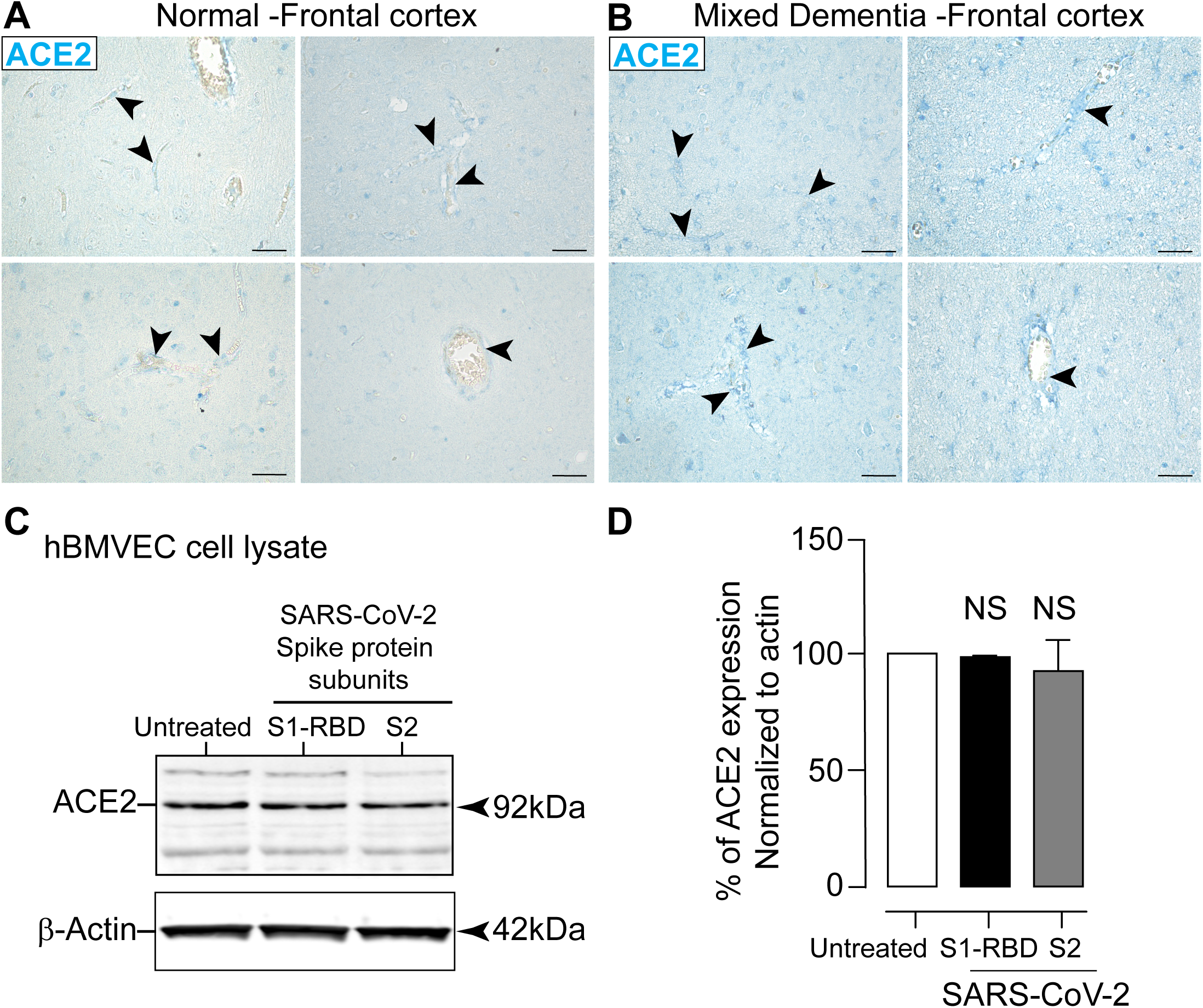
ACE2 is expressed on the cerebral vasculature and in primary human brain microvascular endothelial cells (hBMVECs). Paraffin-embedded brain tissue was sectioned at 5 microns and immunostained for ACE2. (A) shows frontal cortex from a patient with no abnormal neuropathology (control) and (B) from a patient diagnosed with mixed dementia. The representative images at 40x objective magnification, shows the ACE2 expression in blue (Vector Blue). The images were selected to demonstrate immunopositive ACE2 on a range of vessel calibers. Arrow heads indicates the vascular presentation of ACE2 expression compared to ACE2 expression in parenchymal cells. Scalebars = 25 microns. (C) Western blots of hBMVEC cell lysates probed with ACE2 antibodies after cells were exposed to 10nM of SARS-CoV-2 spike proteins S1-RBD and S2. (D) Bar graph of densitometry quantification (average ± SEM) for the expression of ACE2 normalized to β-Actin is shown. Statistical significance differences, *P < 0.05, compared with the untreated control (n=3 donors performed in triplicate).

Currently there is no information that can be found regarding the ACE2 expression profile in primary human microvascular endothelial cells or hBMVECs. Therefore, to determine whether hBMVECs express ACE2 in culture as can be seen in human brain tissue section (Figure 1), western blot analysis was performed on hBMVECs cell lysates. As seen in the Figure 1C, ACE2 is expressed in mature (i.e. postmitotic and with barrier properties) hBMVEC cultures. Furthermore, the expression was unchanged when the cells were exposed to SARS-CoV-2 spike protein subunits S1-RBD (RBD is the binding site on S1 for ACE2) and S2 at 10nM for 24hrs. Densitometry analysis (Figure 1D) of ACE2 bands from the treated cells did not show any difference from untreated controls (average percentage and SEM of 98.5%±0.7 and 92.5%±9.5 respectively).

These results corroborate previous findings on ACE2 detection in CNS tissue and further shows that ACE2 is present in all vascular sized vessels. In addition, ACE2 expression is enhanced in conditions of neuroinflammation. ACE2 expression is also observed in hBMVEC cell cultures and does not change upon exposure of either of the two SARS-CoV-2 spike protein subunits.

### COVID-19 Spike proteins do not affect brain endothelial cell viability

It is possible that the observed vascular effects from SARS-CoV-2 may be explained by cytotoxicity induced by the spike protein. Moreover, to date, no published information exists regarding whether SARS-CoV-2 spike proteins impacts viability of human endothelial cells. Therefore, to assess the cell viability, hBMVECs were exposed to two different concentrations (1nM and 10nM) of S1, S1-RBD (a truncated S1 with only the binding site on S1 for ACE2) or S2 for either 48 hrs or 72 hrs and analyzed using a Live/Dead Cytotoxicity assay. The assay utilizes the combination of live cell-permeable acetomethoxy-derivative of calcein (calcein-AM) and live cell-impermeable ethidium homodimer-1 (EthD-1). The mild detergent, saponin was used as a positive control for inducing cell death. The data is represented as percent of cells that were fluorescently positive at 515 nm (calcein, live cells) or 617 nm (EthD-1, dead cells). The results in Figure 2 shows no evidence of cytotoxicity at 48 hrs with the concentrations tested. Values for live cells were as follows: untreated 99.85% ± 1.04%, saponin 39.35% ±4.09%, S1 1nM 99.78% ± 2.67%, 10nM S1 99.70% ± 1.69%, 1nM S1-RBD 99.49% ± 1.50%, 10nM S1-RBD 99.16% ± 0.92%, 1nM S2 98.65% ± 1.25%, 10nM S2 98.30% ± 2.27%. Values for dead cells were as follows: untreated 0.15% ± 1.04%, saponin 60.65% ± 4.09%, 1nM S1 0.22% ± 2.67%, 10nM S1 0.30% ± 1.69%, 1nM S1-RBD 0.51% ± 1.50%, 10nM S1-RBD 0.84% ± 0.92%, 1nM S2 1.35% ± 1.25%, 10nM S2 1.70% ± 2.27%.

**Figure 2.**
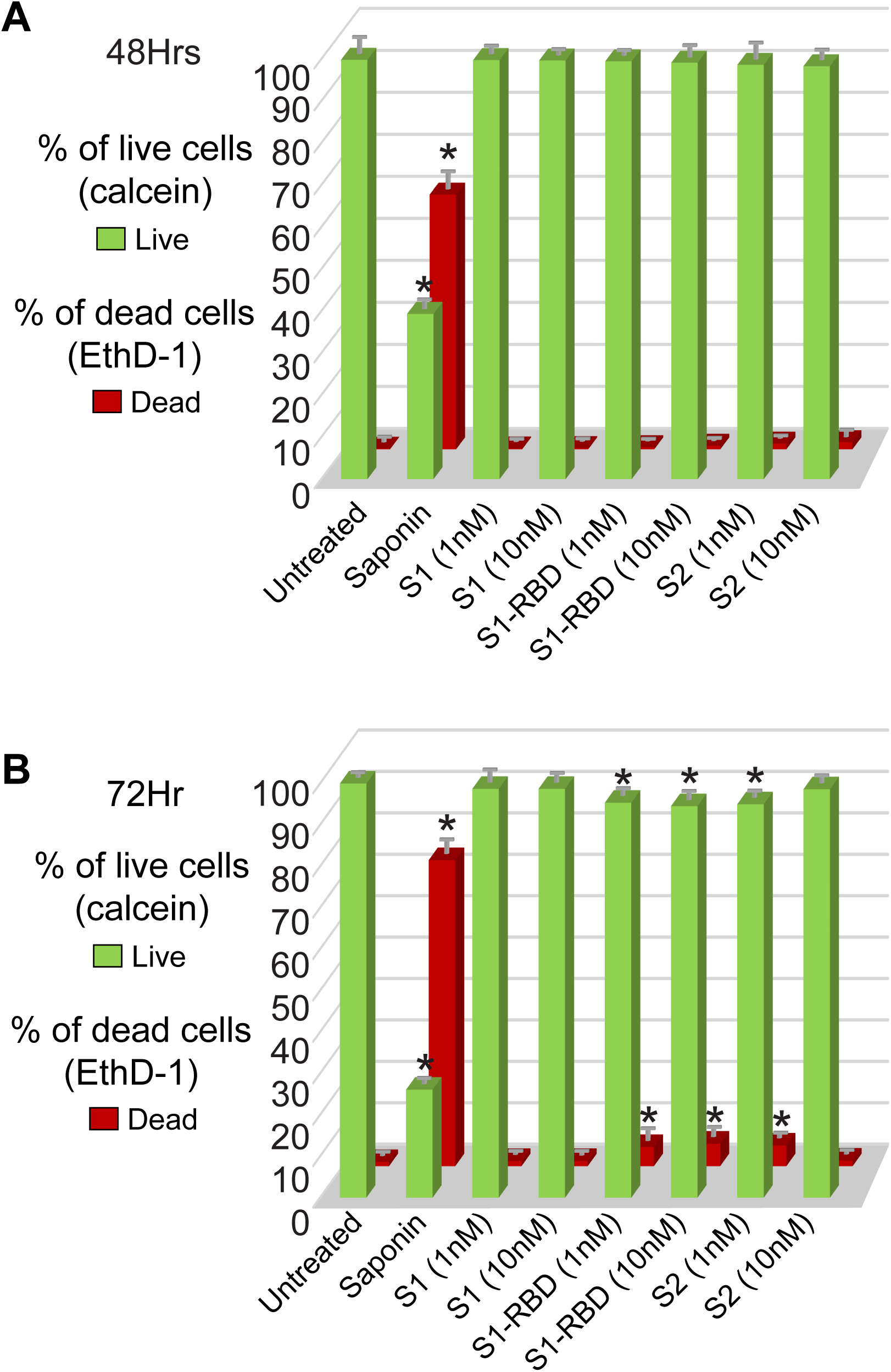
COVID-19 Spike proteins do not affect brain endothelial cell viability. hBMVECs were treated with 1nM and 10nM of the SARS-CoV2 spike proteins (S1 full, S1-RBD, and S2 full) for 48 hrs (A) and 72 hrs (B). Cell viability was determined using the Live/Dead Cytotoxicity assay. Calcein positive (green) indicates live cells while ethidium homodimer-1 (EthD-1, red) indicates dead cells. Saponin was used a positive control. Data obtained from two different donors, each performed in 6 replicates. mean ± SD *p<0.05.

At 72 hrs of incubation with SARS-CoV-2 spike proteins, a slight heightened cell death response with S1-RBD and S2 treated wells was seen. Values for live cells were as follows: untreated 100.08% ± 1.62%, saponin 26.00% ± 1.81%, S1 1nM 98.83% ± 1.19%, 10nM S1 98.83% ± 1.19%, 1nM S1-RBD 95% ± 1.1% (p < 0.05), 10nM S1-RBD 94.58% ± 2.58% (p < 0.05), 1nM S2 94.83% ± 2.41% (p < 0.05), 10nM S2 98.58% ± 3.85%. Values for dead cells were as follows: untreated 0.97% ± 2.0%, saponin 74.00% ± 1.81% (p < 0.001), 1nM S1 1.67% ± 1.19%, 10nM S1 1.67% ± 1.19%, 1nM S1-RBD 4.83% ± 4.41% (p < 0.001), 10nM S1-RBD 5.42% ± 2.58% (p < 0.001), 1nM S2 5.17% ± 2.41% (p < 0.001), 10nM S2 1.42% ± 3.85%.

Overall, these findings indicate that the spike protein subunits from SARS-CoV-2 do not appear to affect brain endothelial cell viability after short-term exposure, suggesting that any pathological effects on the endothelial barrier properties reported in this work is unlikely related to direct cell death by the viral spike protein.

### Loss of BBB integrity induced by the SARS-CoV-2 spike protein

Given that the RBD domain of the S1 SARS-CoV-2 spike protein binds to ACE2 and ACE2 is present in brain endothelial cells, it is possible that engaging the ACE2 protein by the viral spike protein subunits could manifest changes in the status of the barrier. The main feature that distinguishes brain vascular endothelium from endothelium in the periphery is the abundant presence of tight junctions formed between adjacent cells. Tight junctions form the physical barrier of the BBB, preventing the free paracellular flux of ions and small molecules. To this end studies using Electric Cell-Substrate Impedance (ECIS) were performed to determine whether changes to the barrier is observed upon exposure to COVID spike protein. This measure of electrical properties of the endothelial monolayer provides an analytical method for directly evaluating experimental conditions that may induce barrier “tightness”/higher resistance or “leakiness”/lower resistance. ECIS measurements were acquired as described in Materials and Methods. As shown in Figure 3A, the electrical resistance of the monolayers treated with the full S1 subunit of SARS-CoV-2 spike protein reached the lowest and plateaued at 8-12 hrs after the initial exposure (−7.18% ± 2.64% for 10nM, -3.79% ± 1.27% for 1nM) followed by complete recovery in case of 1nM and 10nM S1 and continued decrease in the case of 0.1nM S1-treated cells (down to -10.78% ± 3.52%). In Figure 3B, at 10nM S2, resistance dropped and plateaued earlier (from 6 h to 14 h) and with an average decrease of -7.56% ± 2.43% (p < 0.001) followed by recovery by 24 hrs. 1nM and 0.1nM S2 showed steady gradual decrease throughout whole experiment and reached maximum at 24 hrs (−6.29% ± 1.62% and -6.54% ± 1.75%). In the case of S1-RBD, Figure 3C, evokes a dose-dependent drop of barrier that reached maximum at 14 hrs post-exposure (−12.19% ± 2.74% for 10nM, -8.42% ± 2.53% for 1nM and -5.4% ± 2.84% for 0.1nM RBD).

**Figure 3.**
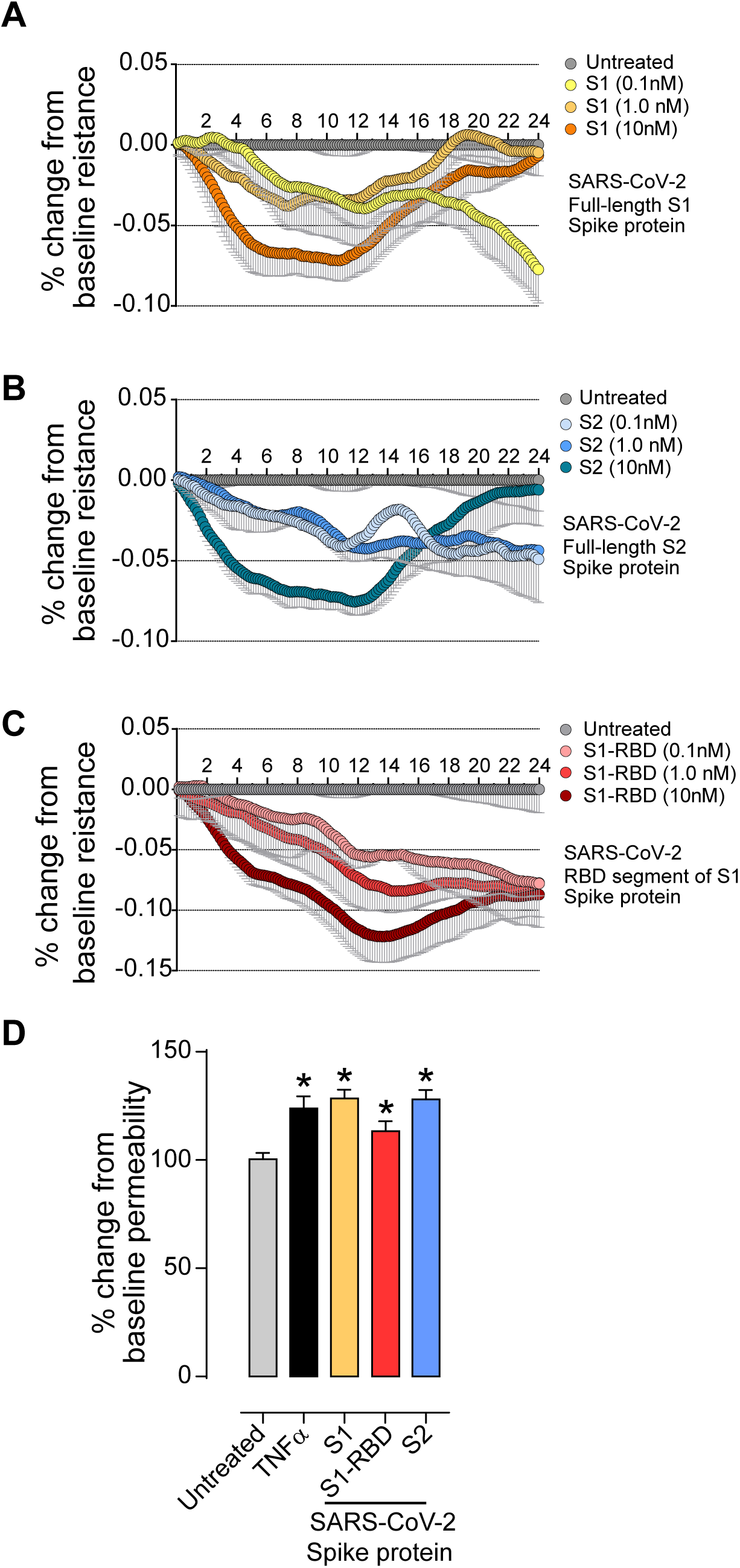
SARS-CoV spike proteins compromise endothelial barrier properties. (A-C) Barrier electrical resistance was modelled based on continuous cell-substrate impedance readings recorded at 6 frequencies (400Hz – 48kHz) every 6 min for the duration of the time shown. Endothelial monolayers were treated with 0.1nM, 1nM or 10nM of S1 full, S1 RBD, S2 full or left untreated to serve as a baseline. Treatments were initiated at 0 timepoint. Experiment was performed in quadruplicates and repeated three times using primary cells obtained from three different donors. Each data point is represented as the percentage of the mean value ± SEM. D. Endothelial monolayers were treated with 100ng/mL TNF-a, 10nM of S1 full, S1 RBD, S2 full or left untreated to serve as a baseline. Barrier permeability to small molecular tracer was modelled using FITC-conjugated DEAE-dextran (3 kDa). Experiment was performed in quadruplicates and repeated three times using primary cells obtained from three different donors. Each data point is represented as the percentage of the mean value ± SD. p-values were computed using one-way ANOVA and Turkey post-hoc test.

To determine how the observed decrease in electrical resistance relates to the flux of molecules across the tight junctions, permeability assays were performed. To focus entirely on paracellular passage via the intercellular junctions we excluded transcytosis and transendothelial channels routes by using smaller molecular weight tracer bearing positive charge (3k Da FITC-conjugated DEAE-dextran). As shown in Figure 3D, at 1 hr, 10ng/mL TNFα (positive control) induced +23.90% ± 5.60% (p < 0.001) permeability, 10nM of S1 subunit induced +28.5% ± 4.11% (p<0.001), 10nM of S2 subunit resulted in +28.1% ± 4.25% (p<0.001), and 10nM of S1-RBD showed no change at 113.3% ± 4.57% (ns).

### Evidence of increased BBB permeability triggered by the SARS-CoV-2 S1 spike protein in a 3D tissue engineered model of the BBB

In order to evaluate the effect of the SARS-CoV-2 spike protein on brain endothelial cells in an in vitro environment that mimics the three-dimensionality of in vivo vasculature, barrier experiments were conducted using a 3D BBB model. The 3D BBB model was perfused for four days with 0.7 dyn/cm^2^ of shear stress to assure formation of tight junctions necessary for barrier function (Figure 4A-G). Following those four days, vessels were perfused for two hours with either 50 nM of S1 spike protein or with untreated medium. Both conditions were exposed to the same magnitude of fluid shear stress, 0.7 dyn/cm^2^, during this period. Assessment of barrier permeability using FITC-dextran indicated a nearly three-fold increase in the permeability coefficient following exposure to the SARS-CoV-2 spike protein for 2hrs (Figure 4F and 4G). Images of the vessels following dextran perfusion validated the permeability measurements; untreated vessels exhibited a sharp gradient of fluorescence at the vessel wall while treated vessels showed substantial leakage (Figure 4F). Moreover, immunostaining for zonula-occludens-1 (ZO-1), a scaffolding protein in tight junction complex, presented with reduced localization of ZO-1 to cell-cell junctions, which is indicative of barrier breakdown (Figure 4D-E). Furthermore, the vessels treated with the viral spike protein featured substantial actin stress fibers within the endothelium, which is characteristic of increased actomyosin contractility and force generation (data not shown). Overall, these results support the findings of the 2D studies by demonstrating spike protein-mediated barrier breakdown in a perfusable 3D configuration.

**Figure 4.**
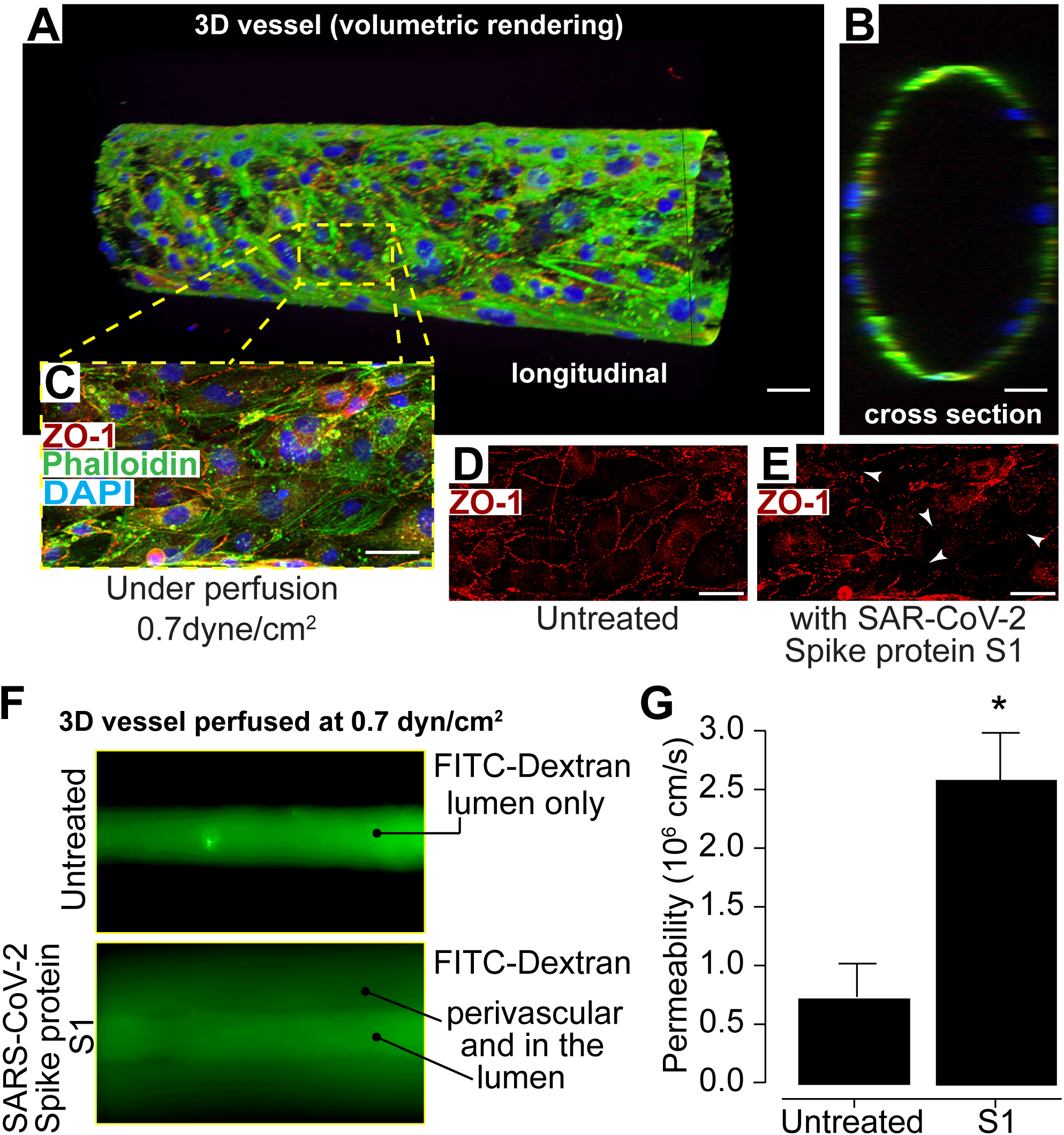
Viral S1 spike protein alters barrier status in a 3D tissue engineered microfluidic model of the human BBB. Confocal microscopy and volumetric rendering was used to visualize tissue engineered vessel. (A) Shows a longitudinal view of an endothelialized void after perfusion that formed a predictive vessel geometry analogous to those found within the brain. (B) provides a cross sectional perspective indicating a single layer of endothelial cells. In (C) a representative merged image of engineered vessel constructs fixed and immunestained for the tight junction protein, ZO-1, along with phalloidin to label actin and also the nuclear stain, DAPI. (D) shows the typical ZO-1 membranous pattern expected in mature barrier forming brain endothelial cells. (E) after perfusion for 2 hrs of SARS-CoV-2 spike protein subunit 1 (10nM), constructs were also fixed and immunolabeled for ZO-1. The arrows point to areas in which the ZO-1 cellular pattern is discontinuous, punctate or absent signifying areas of barrier breach. Scalebar = 20 microns. (F) Fluorescence intensity after ten minutes of perfusion with 4 kDa FITC-dextran, indicating the impaired barrier function in vessels perfused after 2 hrs of the S1 spike protein versus untreated controls. G) Quantitative measurements for permeability coefficients of vessels exposed to the SARS-CoV-2 S1 spike protein subunit compared to untreated controls. n=3. p<0.05.

### The SARS-CoV-2 spike protein triggers brain endothelial activation

The experiments above showed that SARS-CoV-2 spike protein significantly affects barrier integrity. It is possible that the viral spike protein triggers the activation and the pro-inflammatory response of the endothelial cells which results in barrier dysfunction. To test this possibility, flow cytometry experiments were performed to analyze the surface expression an of intracellular adhesion molecule-1 (ICAM1) and vascular cell adhesion protein-1 (VCAM1) expression as a function of time (4hr, 24hr). S1, S1-RBD and S2 elicited a robust increase in ICAM-1 (Figure 5A-C,E) and VCAM-1 (Figure 5F-H,J) by 4hrs and which remained elevated at 24hrs. TNF-α was used as a control for endothelial activation (Figure 5D,E,I,J). Baseline mean fluorescent intensity (MFI) was 565.7 ± 19.1 for ICAM-1 and 394.0 ± 19.2 for VCAM-1. The full S1 subunit elevated the MFI for ICAM-1 to 2279.0 ± 60.7 (4 hrs) and 2644.7 ± 93.1 at 24hr and represented a 4-4.5-fold increase in MFI (Figure 5A, E). VCAM MFI was similarly affected and increased to 1281.3 ± 28.5 (4 hrs) and 1572.7 ± 43.3 (24hr) (Figure 5F,J). The response to the S1-RBD truncated form also increased ICAM-1 and VCAM-1, albeit to a lesser degree. ICAM-1 MFI was 1223.0 ± 49.7 (4hr) and 1632.0 ± 160.1 (24hr) (Figure5B, E) while VCAM-1 MFI increased to 718 ± 28.1 (4hr) and 1101.3 ± 88.5 (24hr) (Figure 5G,J). Finally, treatment with S2 full induced a similar response to S1 full with ICAM-MFI increasing to 2203.6 ± 61.3 at 4hr and 2409 ± 154.7 at 24 hrs (Figure 5C,E). Similarly, VCAM-1 MFI increased to 1340.0 ± 56.7 at 4hr and 1487.3 ± 74.6 at 24 hrs (Figure 5H, J). While the increase in ICAM-1 and VCAM-1 MFI was to a similar magnitude by 4 hrs for the S1 full and S2 full compared to TNF-α, the response to the spike proteins plateaued and did not increase further by 24 hrs as was seen with TNF-α.

**Figure 5.**
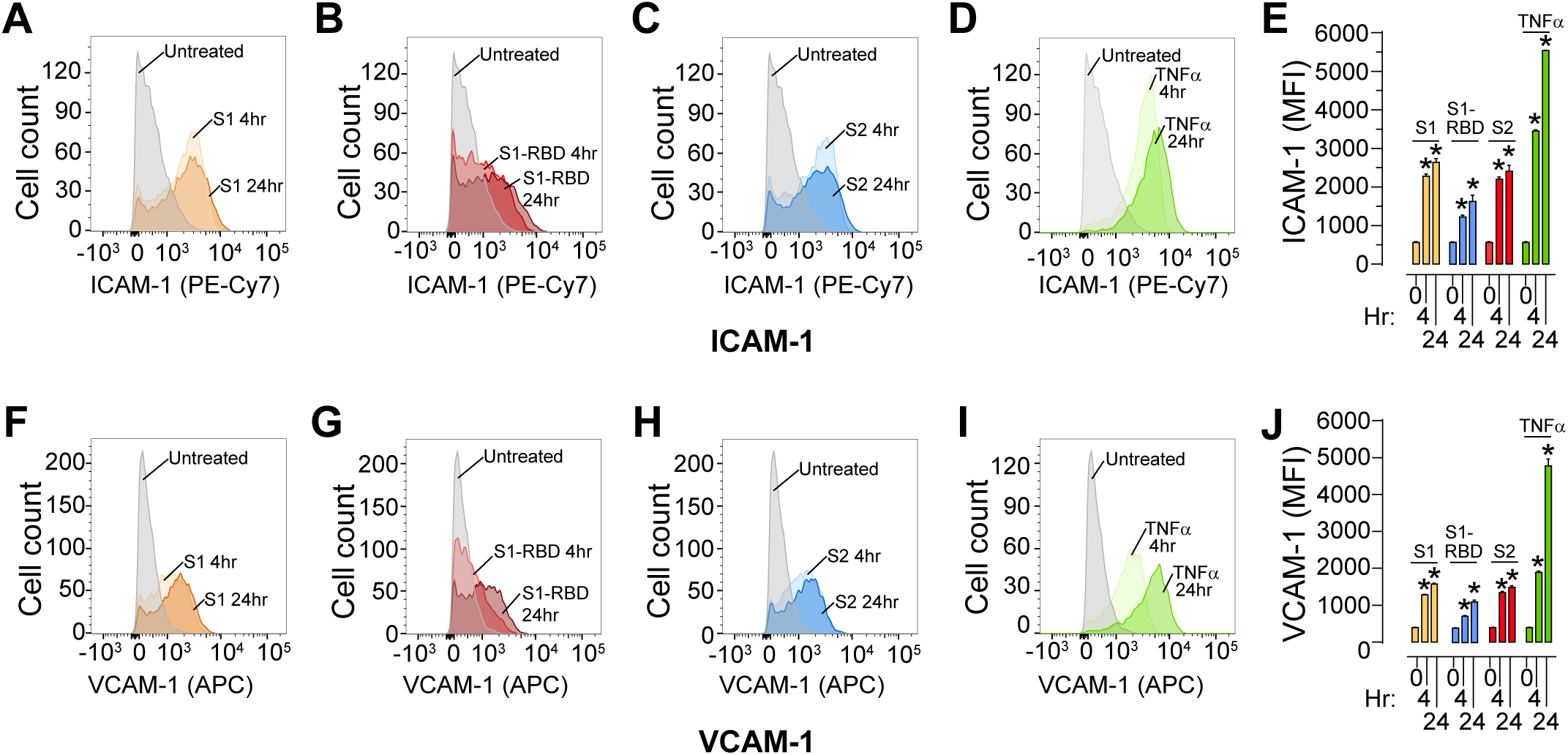
SARS-CoV-2 spike proteins triggers the enhanced surface expression of adhesion molecules. Human brain microvascular endothelial cells (hBMVECs) were treated with 10nM of S1 full, S1 RBD, S2 full or 100ng/mL of TNF-α for 4 and 24hr. Cells were stained for ICAM-1 and VCAM-1 expression and analyzed using a FACS Canto II flow cytometer. Shown are representative histograms for ICAM-1 expression in response to S1 full (A), S1-RBD (B), S2 full (C), TNF-α (D) and the bar graph quantification of the Mean Fluorescent Intensity (MFI) (E). Representative histogram for VCAM-1 expression in response to S1 full (F), S1 RBD (G), S2 full (H), TNF-α (J) and the bar graph quantification of MFI (J). (n=3, *p<0.05).

Together, these results provide evidence for induction of a pro-inflammatory phenotype when hBMVECs are exposed to the two subunits of the SARS-COV-2 spike protein

### SARS-CoV-2 viral protein triggers a pro-inflammatory response and upregulation of MMPs in BMVECs

One possibility that could offer an explanation to the destabilizing effects seen on barrier function by the spike protein may be due to further inflammatory response, in this case gene expression of cytokines. Thus, the inflammatory response of BMVECs expose to the various SARS-CoV-2 spike protein subunits was assessed using gene expression assays and analyzed to the baseline (untreated) expression. As shown in Figure 6A-B, IL-1β gene expression was statistically upregulated for cells incubated in the presence of S1-RBD (4 hrs treatment: 2.313 ± 1.05, p < 0.001) and S1 (4 hrs treatment: 2.378 ± 0.865, p < 0.001 and 24 hrs treatment: 2.412 ± 0.317, p < 0.001). Similarly, IL-6 mRNA was significantly upregulated after 4hrs of S1 (1.796 ± 0.533, p = 0.014) S1-RBD (2.383 ± 1.311, p < 0.001) and S2 (2.051 ± 0.449, p < 0.001). While 24 hrs treatment yield significant results only for S1 (1.779 ± 0.702, p = 0.017) A marked significance in CCL5 gene expression was seen in all conditions tested and at both time points (S1 at 4hrs 9.793 ± 7.151, p < 0.0001 and at 24hrs 13.99 ± 5.093, p < 0.001; S1-RBD at 4hrs 6.181 ± 1.852, p = 0.026 and at 24hrs 6.344 ± 1.603, p = 0.02; S2 at 4hrs 6.644 ± 4.352, p = 0.01 and at 24hrs 8.529 ± 5.959, p < 0.005). CXCL10 mRNA was significantly upregulated in S1-treated cells at 4 hrs (16.62 ± 7.33, p < 0.001) and at 24 hrs (5.251 ± 1.686, p = 0.014); and after 4 hrs of S2 treatment (6.963 ± 4.409, p < 0.001). Although inflammatory responses by brain endothelial cells is counteractive to barrier stability. It is possible that other cellular process may contribute to the observed increased in barrier permeability. Another likely molecular target known to induce a compromise to barrier integrity is the family of Matrix metalloproteinases (MMPs). Thus, the gene expression of MMPs was examined as another mechanism for which SARS-CoV-2 spike proteins may compromise barrier maintenance. As shown in Figure 6C-D, 4hrs of S1-RBD exposure to BMVECs resulted in elevated gene expression (fold change ± SD) of MMP2 (1.406 ± 0.251, p = 0.012), MMP3 (4.398 ± 0.774, p = 0.003), MMP9 (1.720 ± 0.485, p < 0.005), and MMP12 (3.271 ± 1.558, p < 0.001). After 24hrs of S1-RBD exposure, MMP2 (1.378 ± 0.446, p < 0.022), MMP3 (4.319 ± 1.45, p = 0.0004), MMP9 (1.66 ± 0.578, p = 0.011) and MMP12 (3.01 ± 0.612, p < 0.001) were all upregulated. S1 exposure at 4 hrs affects only MMP3 (3.401 ± 1.01, p = 0.016) and MMP12 (3.416 ± 1.405, p < 0.001) gene expression. However, at 24hrs of S1 exposure results in elevated MMP3 (5.557 ± 1.763, p < 0.0001), MMP9 (1.594 ± 0.879, p = 0.027), and MMP12 (3.305 ± 0.759p < 0.0001). Similarly, S2 at 4hrs induced upregulation of MMP3 (7.849 ± 3.715, p < 0.0001) and MMP12 (3.021 ±0.446, p < 0.0001), with unchanged MMP2 and MMP9. After 24h of S2 exposure, MMP2 (1.613 ± 0.173, p < 0.001), MMP3 (3.520 ± 1.751, p = 0.010), MMP9 (1.705 ± 0.340, p = 0.0058), and MMP12 (4.076 ± 1.023, p < 0.001) were all significantly different from untreated samples.

**Figure 6.**
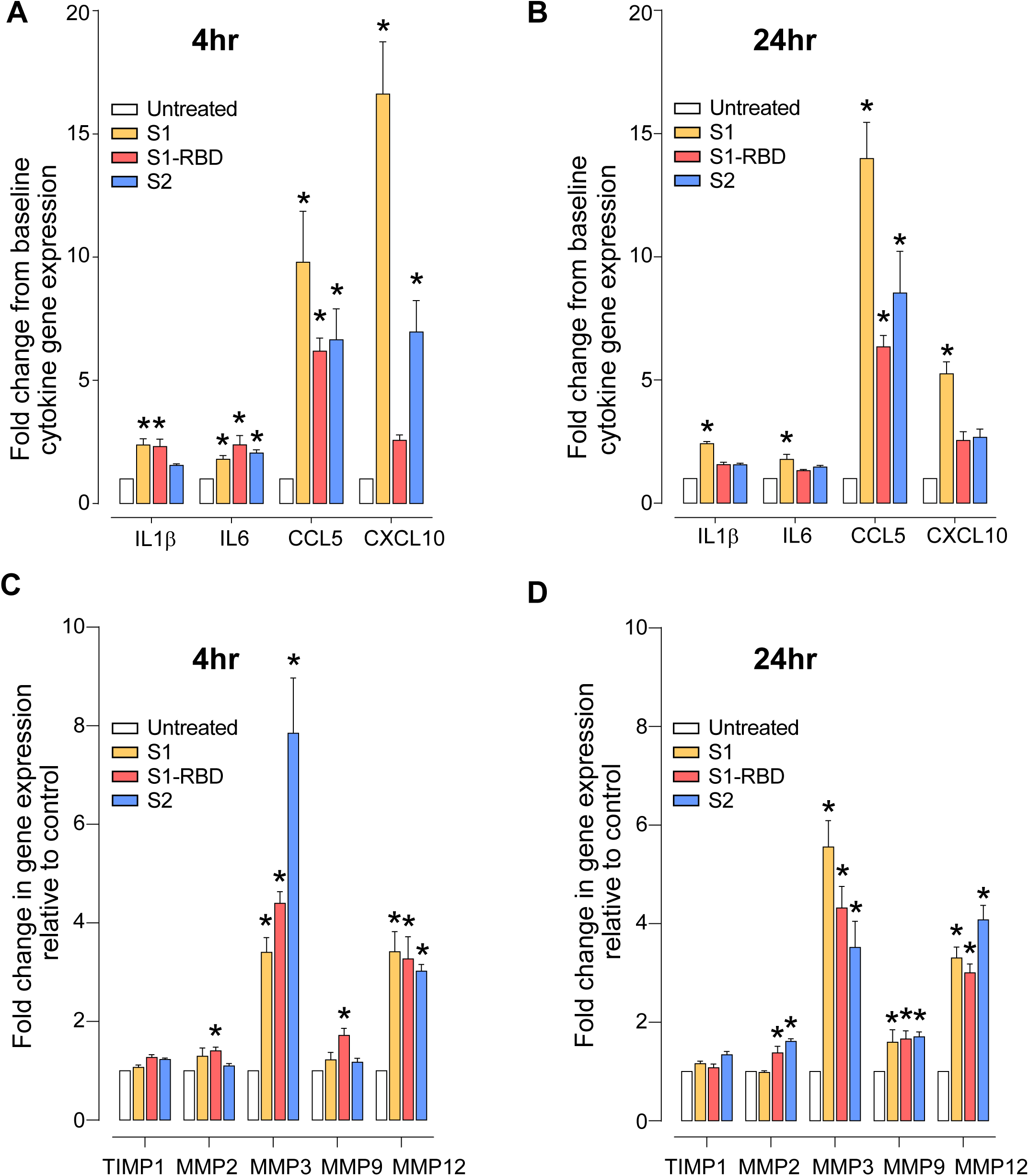
SARS-CoV-2 viral proteins trigger pro-inflammatory responses and upregulation of MMPs in BMVECs. Confluent BMVECs monolayers were incubated with 10nM of S1 full, S1-RBD, S2 full for time indicated or left untreated to serve as a control. mRNA expression of cytokine genes after stimulation with the SARS-CoV-2 spike protein S1, S1-RBD and S2 subunits (all at 10nM) are shown for either 4 hrs and 24 hrs. Taqman gene expression assays were performed in quadruplicates and repeated three times using primary cells obtained from three different donors. Target cytokine genes analyzed included: IL1β, IL6, CCL5, CXCL10 at 4hrs (A) and 24 hrs (B). Gene expression analysis for MMP2, MMP3, MMP9, MMP12 and the MMP inhibitor TIMP1 are shown for 4 hrs (C) and 24 hrs (D) respectively. Each bar represents fold-change ± SEM. Data sets were separately analyzed using one-way ANOVA and p-values were computed using Turkey post-hoc test.

These analysis supports the notion that SARS-CoV-2 spike proteins can trigger a very specific pro-inflammatory response (that includes MMP expression) in brain endothelial cells thereby offering a means for which SARS-CoV-2 may breach the BBB.

## DISCUSSION

The emergence of the COVID-19 pandemic (as a result of infection from SARS-CoV-2) and its ensuing consequences on public health has dramatically changed our way of life. Those with COVID-19 can be asymptomatic or present with a wide array of symptoms which critically influences who recovers from the infection. In more severe cases, patients can progress to acute respiratory distress syndrome, septic shock, metabolic acidosis, coagulopathy and multiple organ dysfunction. The host’s pro-inflammatory response, particularly in cases of aggressive inflammatory phenotypes, strongly contributes to disease prognosis. Neurological observations as nausea, headache, anosmia, myalgia, impaired consciousness, and acute cerebrovascular diseases have also been reported in some COVID-19 patients^17,35,36^. It is clear that much remains unknown about the pathophysiology of SARS-CoV-2 infection not only in periphery organ systems but specially in regard to the CNS.

The analysis within this report provides evidence that the SARS-CoV-2 spike protein can directly affect the status of the blood-brain barrier (BBB) which gives insight into the neuropathology associated with COVID-19. In particular, the breakdown of the (BBB) by this novel coronavirus offers a possible avenue for counteracting the consequences of acute ischemic stroke observed in COVID-19 patients younger than 50 years old^37^.

COVID-19 can induce microclots in both the vasculature of periphery tissues but also within vessels of the CNS. In fact, Bryce et al. found that 6/20 cases had microthrombi and acute infarction in the brain^31^. Therefore, future studies should place focus on interrogating the connection between virus-mediated barrier disruption and coagulation to determine the mechanisms unique to the cerebral vasculature responsible for heightening the risk of strokes in COVID-19 patients.

Angiotensin converting enzyme 2 or ACE2 is the primary membranous cellular binding target for the SARS-CoV-2 spike protein. ACE2 expression has been detected previously in the brain vasculature^19^ and recently confirmed in a study of COVID-19 patients by Bryce et al^31^. Whether ACE2 expression changes in patients with comorbidities (hypertension, diabetes etc.) is unknown. However, it is known that in cases of neurodegeneration, Kehoe et al.^19^ found that ACE2 activity in brain tissues from Alzheimer’s (AD) patients was reduced. Unfortunately, the above study did not provide a comparison of protein expression with control cases. The results herein make two important assertions in comparisons between normal cortex vs. cortex from mix dementia cases. The first is that ACE2 expression can be found in a wide range of vessel calibers that includes capillaries, arterioles, and venules (Figure 1). The second, is that ACE2 expression appears upregulated in the parenchyma and in larger vessels of cases of mixed dementia. This expression suggests that SARS-CoV-2 could encounter its key binding target in the cerebrovasculature. Interestingly in Bryce et al., virus like particles were readily detectable by electron microscopy in the frontal lobe endothelium of a COVID-19 patient^38^.

It is now well accepted that COVID-19 can strike all age groups, including children. An observed complication of COVID-19 infection in some children is the emergence of severe Kawasaki-like disease characterized by multisystemic inflammation^39^. A hallmark feature of Kawasaki disease is vasculitis that results from the inflammation generated by affected endothelium. Interestingly much of COVID-19 systemic pathogenesis can be explained by the virus impact on the endothelium^30^.

The BBB is the interphase that is breached by various neuroinvasive viral pathogens including rabies^40,41^, HIV-1^42-44^, West Nile^45,46^, Zika^47^, and influenza^48^. Viruses negatively affect the BBB not only by direct interaction with endothelial cells but also by induction of host immune responses that result in elevated expression of pro-inflammatory cytokines, chemokines, cell adhesion molecules that lead to a demise in BBB structural and functional integrity^49^. A disruption in the BBB facilitates crossing of viral particles and infected immune cells further elevating levels of inflammatory mediators^49-52^. SARS-CoV-2 could employ such mechanisms of neuroinvasion and future studies will help discern whether this is the case. However, during the course of any viral infection the shedding of viral proteins are produced, as such the effect of the essential SARS-CoV-2 spike protein was evaluated for its ability to induce dysfunction of the BBB. First, to rule out the possibility of a cytotoxic effect by SARS-CoV-2 spike protein (S1 and S2 subunits) on endothelial cells, live/dead assays were performed (Figure 2). It was found that at early timepoints up to 48 hrs, there was no considerable cell death caused by either the full length S1, the S1 truncated form containing the RBD or S2. At the 72 hrs time point, the results showed some increased in cell death that reached statistical significance. Thus, it appears that by chronic (longer that 72 hours) exposure to SARS-CoV-2 spike proteins does induce cell death of brain endothelial cells that actually corroborates with recent clinical data^30^.

Next, the effect of SARS-CoV-2 spike proteins were evaluated for their ability to cause a loss in brain endothelial barrier function. To evaluate barrier integrity, experiments were performed that measured the electrical resistance (an analytical means to examine barrier “tightness”) and paracellular permeability of hBMVECs exposed to different subunits of SARS-CoV-2 spike protein. Even single application of the S1, S1-RBD or S2 to endothelial monolayers resulted in a dose-dependent loss of the barrier electrical resistance that peaked at 12 – 14 hrs. Interestingly, high doses of full length S1 and S2 subunits both caused transient loss of electrical resistance that was completely recovered by 24 hrs, raising the possibility that structural reorganization at the tight junction complex occurs rather than outright loss of tight junctions (Figure 3). On the other hand, the truncated S1 form with only the RBD sequence caused irreversible loss of barrier function (Figure 3C) that plateaued at 24 hrs. These results indicate that ACE2 receptors may not be the exclusive point of contact between SARS-CoV-2 and brain endothelial cells. Most likely the interaction between SARS-CoV-2 spike proteins and the BBB is multifocal and involves reversible activation at more than one receptor or signaling cascade. To determine whether the decrease in transendothelial resistance by the spike proteins result in a hyper-permeable barrier, permeability assays were subsequently performed. The result in Figure 3D shows that SARS-CoV-2 spike protein significantly increased the rate of passive paracellular passage of small molecular tracers providing a second indicator of barrier dysfunction.

Endothelial cells are configured as vessels and are constantly exposed to fluid shear stress from the blood. Therefore, it is important to validate findings from static systems in BBB models that factor physiologic parameters such as dynamic flow and intercellular geometries. To this end, experiments were performed using brain endothelial cells grown in a cylindrical space within a gel matrix as described previously^33^. After endothelialization, perfusion was introduced to promote barriergenesis that generates properties featured at the BBB. In Figure 4 these vascular constructs are shown to mature as a single layer (cross section) of endothelium that form intercellular tight junctions and restrict movement of fluorescent tracers. These systems (when also coupled with other cells) represent the most advanced recapitulation of the human neurovascular unit. Once SARS-CoV-2 spike protein subunit S1 was introduced, the presence of barrier permeability (from lumen to parenchymal compartment) was clearly evident as early as 2 hrs. These results suggest that whether free viral spike proteins or those on the surface of the virus present during COVID-19 infection could induce barrier permeability (albeit once a certain threshold is reached) equivocal to the concentrations used here. As this is the first report on the topic, much work remains, particularly in regard to how permeability dynamics may change once these 3D microfluidic constructs are used with the whole SARS-CoV-2 virus.

Endothelial cells are an essential part of the inflammatory response since activation of the endothelium allows for recruitment and mobilization of immune cells to the tissues that are under pathogen attack. Once activated, brain endothelial cells upregulate expression of cell adhesion molecules (CAMs) and pro-inflammatory cytokines that play a key initial role in the process of neuroinflammation. Cell adhesion molecules ICAM-1 and VCAM-1 are key to the transendothelial migration of immune cells in response inflammatory challenge and are often upregulated in viral infection^53,54^. SARS-CoV-2 spike proteins (S1, S2), like other virus proteins, activated hBMVECs, upregulating surface ICAM-1 and VCAM-1 (Figure 5) and in conjunction with reduced barrier tightness (Figure 2). This indicates the potential for enhanced immune infiltration into the CNS and possible neuroinvasion.

Another measure of cellular activation is the upregulation of cytokine genes. Cell exposed to the spike viral protein showed gene expression increase in the leukocyte chemotaxis factor, CXCL10 and CCL5 (RANTES) (Figure 6). Of note, mRNAs for interleukins, IL-1β and IL-6, were significantly altered in response to spike proteins. Endothelial activation also features increased expression of matrix metalloproteinase or MMPs, a family of enzymes involved in the remodeling of extracellular matrix in both normal physiological and pathological processes. Activated by pro-inflammatory cytokines^55^, MMPs also regulate tight junction proteins degradation and post-translational modifications^56-58^. In this study, we report that the spike protein increases MMP3 and MMP12, and to a lesser extent MMP2 and MMP9 mRNA expression. MMP3 has been previously implicated in traumatic brain injury^59^ by digesting tight junctions proteins followed by the BBB opening^60^. These reports corroborate with our findings of decreased barrier resistance (Figure 3) and heightened secretion of chemotactic chemokines (Figure 5). MMP12, on contrary, is not involved in the BBB damage, but plays role in immune cells extravasation and migration into the brain^61^. Taking together our data of elevated MMP3, CCL5, CXCL10 and CAMs gene expression, we can speculate that SARS-CoV-2 is potentially neuroinvasive virus as it turns on the machinery to facilitate the migration of infected immune cells “Trojan horses” into the brain parenchyma.

To our knowledge, this is the first reported evaluation that examined the effects of SARS-CoV-2 spike protein on the blood brain barrier. Our findings provide insight into the continued theme that this novel coronavirus triggers specific responses at the endothelium. Specifically, in regards to the brain endothelium, the SARS-CoV-2 spike protein induced destabilization of the BBB, promoted a pro-inflammatory status but did not appear alter cell viability acutely. A dysfunction of the barrier, particularly in terms of where it occurs neuroanatomically, offers an explanation to the possible observed neurological complications seen in COVID-19. Lastly, the opening of the BBB, hints at the possible means in which the SARS-CoV-2 pathogen could also neuroinvade.

## ACKNOWLEDGEMENTS

Methodologies and reagents used were developed in part by the following funding sources: T32 DA007237 (TPB), 5R01DA046833 (SHR & RP) and K01DA046308 (AMA). Human fetal tissue for the isolation of human fetal brain microvascular endothelial cells was obtained from the Birth Defects Research Laboratory which is supported by NIH award number 5R24HD000836 from the Eunice Kennedy Shriver National Institute of Child Health and Human Development. We would like to recognize the Temple University IBC committee (particularly program coordinator, Mary B. Pultro) for their attentive and efficient review which allowed these experiments to be performed in a timely manner. The authors declare no competing financial interests.

